# Future spinal reflex is embedded in primary motor cortex output

**DOI:** 10.1101/2023.06.19.545500

**Authors:** Tatsuya Umeda, Osamu Yokoyama, Michiaki Suzuki, Miki Kaneshige, Tadashi Isa, Yukio Nishimura

**Affiliations:** Department of Developmental Physiology, National Institute for Physiological Sciences, National Institute of Natural Sciences, Okazaki, Aichi, 4448585, Japan; Department of Integrated Neuroanatomy and Neuroimaging, Graduate School of Medicine, Kyoto University, Kyoto, 6068501, Japan; Department of Neurophysiology, National Center of Neurology and Psychiatry, Kodaira, Tokyo, 1878502, Japan; Neural Prosthetics Project, Tokyo Metropolitan Institute of Medical Science, Setagaya, Tokyo, 1568506, Japan; Department of Neuroscience, Graduate School of Medicine, Kyoto University, Kyoto, 6068501, Japan; Human Brain Research Center, Graduate School of Medicine, Kyoto University, Kyoto, 6068507, Japan; Institute for the Advanced Study of Human Biology (WPI-ASHBi), Kyoto University, Kyoto, 6068510, Japan; School of Life Science, The Graduate University for Advanced Studies (SOKENDAI), Hayama, Kanagawa, 2400193, Japan; PRESTO, Japan Science and Technology Agency (JST), Kawaguchi, Saitama, 3320012, Japan

## Abstract

Mammals can execute intended limb movements despite the fact that spinal reflexes involuntarily modulate muscle activity. To generate appropriate muscle activity, the cortical descending motor output must coordinate with spinal reflexes, yet the underlying neural mechanism remains unclear. We simultaneously recorded activities in motor-related cortical areas, afferent neurons, and forelimb muscles of monkeys performing reaching movements. Motor-related cortical areas, primarily the primary motor cortex (M1), encode subsequent afferent activities attributed to forelimb movements. M1 also encodes a subcomponent of muscle activity evoked by these afferent activities, corresponding to spinal reflexes. Furthermore, selective disruption of the afferent pathway specifically reduced this subcomponent of muscle activity, suggesting that M1 output drives muscle activity not only through direct descending pathways but also through the “transafferent” pathway composed of descending plus subsequent spinal reflex pathways. Thus, M1 provides optimal motor output based on an internal forward model that prospectively computes future spinal reflexes.

## Introduction

The organization of our motor system is characterized by a nested hierarchy comprising supraspinal and spinal structures. To conduct voluntary limb movements, the primary motor cortex (M1) sends a motor command through descending pathways to the spinal cord to generate muscle activity ^1, 2^. In addition, the spinal reflex modulates muscle activity by transforming somatosensory signals from peripheral afferents during voluntary movements ^3-6^. Inputs from both descending motor and afferent somatosensory pathways ultimately converge on spinal motor neurons as a final common pathway ^7-9^. The temporally organized integration of cortical motor commands and somatosensory feedback signals in spinal motor neurons generates the appropriate muscle activity needed to execute an intended movement ^10, 11^. Therefore, a coordinated interplay between the descending motor drive and somatosensory feedback signals has a pivotal role in the generation of muscle activity during voluntary movements ^12^. However, the mechanism by which the descending motor drive coordinates with spinal reflexes to generate appropriate muscle activity remains unclear. Here, we study the flow of neural information across the motor-related cortical areas (MCx), peripheral afferents, and limb muscles in voluntary forelimb movements to provide evidence for the incorporation of information from spinal reflexes driven by somatosensory afferent activity into descending control signals from the M1.

## Results

### Multiregional recording in limb movement

To explore a coordinated interplay between descending motor drive and somatosensory feedback signals in voluntary limb movements, we conducted the simultaneous recording of electrocorticographic (ECoG) signals from the MCx, including the M1 and dorsal (PMd) and ventral (PMv) premotor cortices, the activity of a population of peripheral afferents at the lower cervical level (25-39 units from C7 and C8 dorsal root ganglia (DRGs) of monkey T, and 11-15 units from C6 and C7 DRGs of monkey C), electromyographic (EMG) signals from the forelimb muscles (12 and 10 muscles from monkeys T and C, respectively), and kinematics of the forelimb joints (wrist, elbow, and shoulder) in two monkeys, as the monkeys performed reaching and grasping movements (Fig. 1a,b and Extended Data Fig. 1) ^10, 13^. An example of simultaneous multiregional recording data (monkey T, three trials) is illustrated in Fig. 1b. It is widely accepted that cortical high-gamma activity reflects the activity of the neuronal population below the electrode; thus, we analyzed high-gamma activity (60-180 Hz) in the MCx ^14^.

**Fig. 1.**
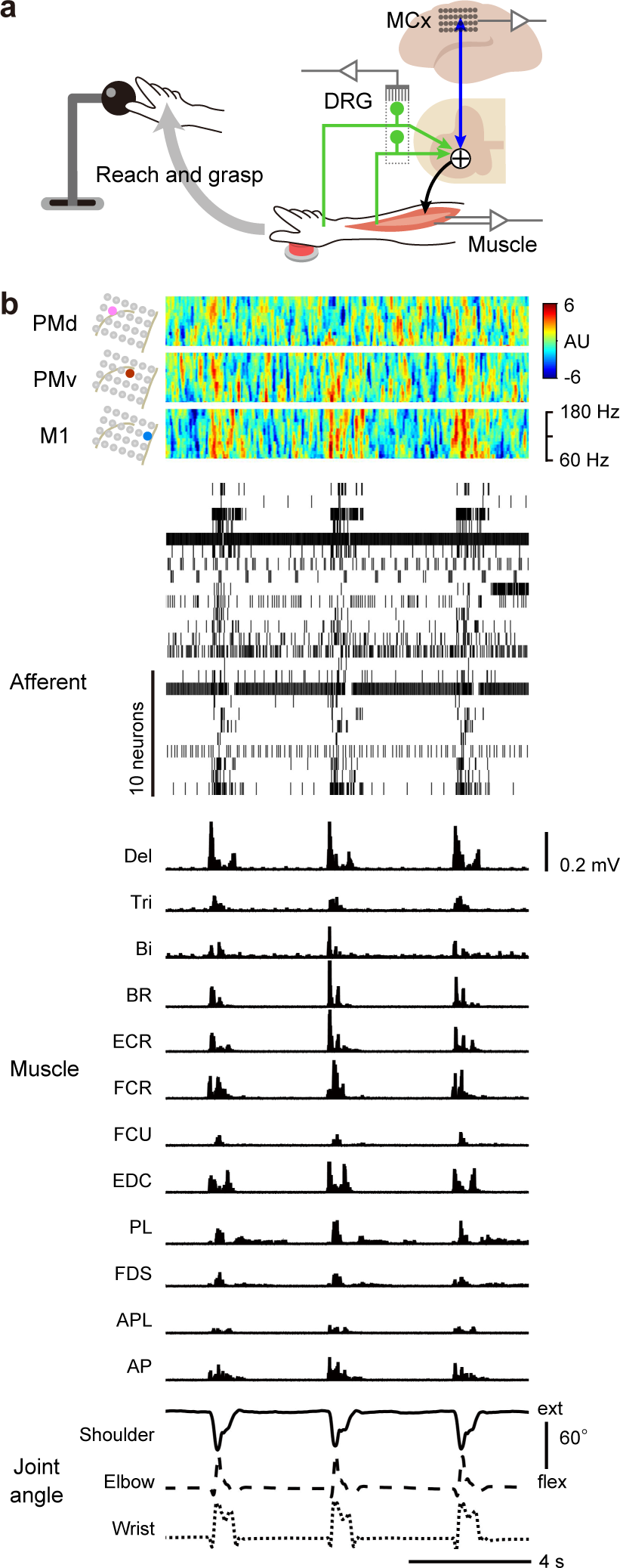
Multiregional recordings during voluntary upper limb movements. **a**, Experimental setup. **b**, Simultaneous recordings in three trials. *Top*: Power spectrograms in the MCx. *Second*: Raster plots of peripheral afferent activity. *Third*: Activity of forelimb muscles. *Bottom*: Forelimb joint angles along the extension-flexion axis.

## MCx encodes reafferent signals

The MCx sends descending control signals to spinal motor neurons to produce muscle activity for the execution of an intended movement ^15^. Muscle activation and subsequent joint movements further lead to the activation of peripheral afferents, which represents reafferent signals ^16^. According to this sequential flow of information, we reasoned that MCx activity is related to the generation of subsequent afferent activity evoked by limb movement (Fig. 2a). It has been well established that muscle activity can be explained by a linear sum of MCx activity that occurs 5–50 ms prior to the muscle activity to be calculated ^13, 17-19^. Therefore, we assumed a delayed linear sum of MCx activity as a first-order model of afferent activity. By constructing a linear model to explain instantaneous afferent activity using MCx activity during the 50 ms preceding the timing of afferent activity, we were able to accurately reconstruct the overall temporal pattern of activity of a substantial number of peripheral afferents; this model outperformed models constructed using shuffled controls (Fig. 2b,c). This result suggests that MCx activity encoded subsequent activity of peripheral afferents.

**Fig. 2.**
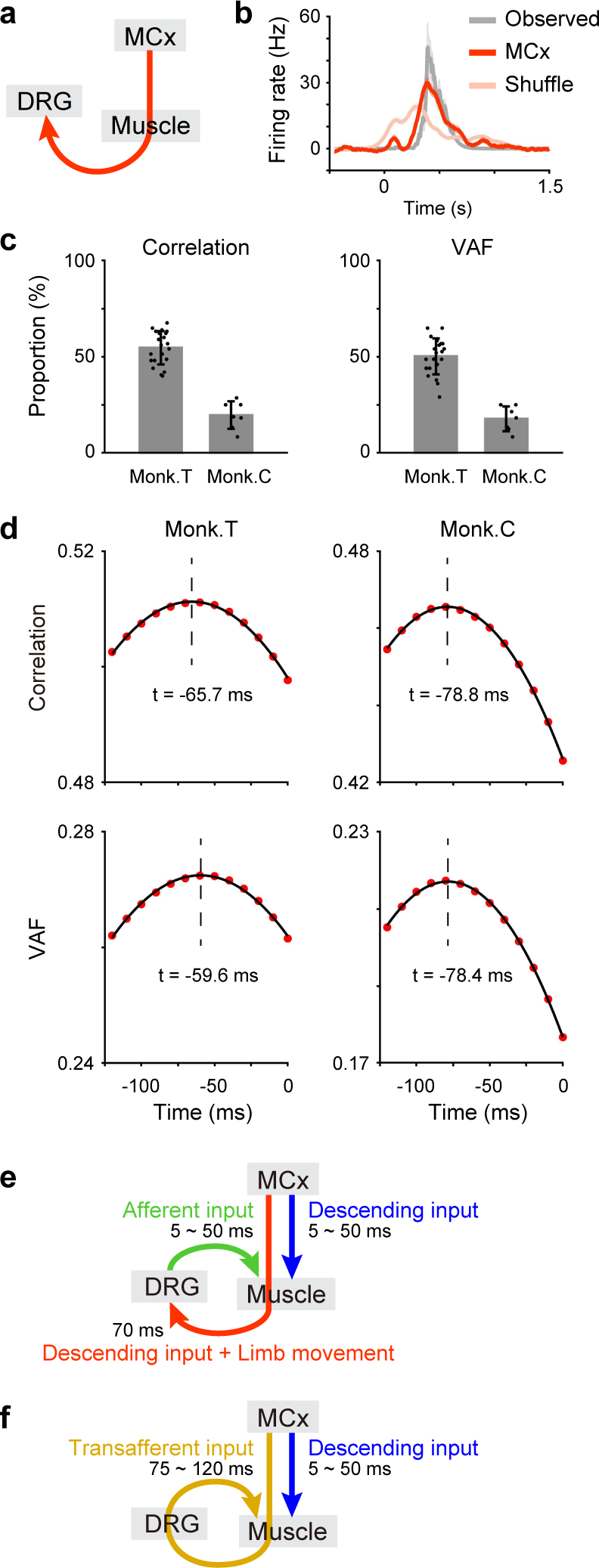
The MCx encodes reafferent signals. **a**, Model accounting for afferent activity in DRGs from MCx activity. **b**, The observed afferent activity, reconstruction using activity in the MCx and shuffled control data aligned to movement onset. The shaded areas indicate the SEM. **c**, Proportion of afferents whose activities were more accurately reconstructed from MCx activity than from the shuffled data in the correlation coefficients and variances accounted for (VAFs) between the observed and reconstructed traces (monkey T, 21 sessions; monkey C, 7 sessions). Data are the mean ± SD. **d**, Correlation coefficients and VAFs plotted against the lag times between afferent and MCx activity. The black line represents the fit to a quadratic curve. The vertical dotted lines indicate the lag time at the maximum value of the fitted curve. **e**, Schematic illustrating a plausible sequence of information flow from the MCx to muscles during voluntary movements. **f**, Same as in **e**, but MCx signals activate muscles via the descending pathway (blue) and via the transafferent pathway (orange).

To determine the timing at which descending input was most associated with subsequent afferent activity, we built models with different lag times between afferent and MCx activity and calculated the reconstruction accuracy of these models. MCx activity prior to approximately 70 ms (monkey T: 62.7 ms, monkey C: 78.6 ms) was found to have an adequate lead time to explain subsequent afferent activity (Fig. 2d). These results suggest that MCx output is transmitted to peripheral afferents over 70 ms (a red arrow in Fig. 2e).

## Spinal reflex pathways convey MCx output

Our recent study revealed that afferent activity associated with forelimb movements contributed significantly to muscle activity (a green arrow in Fig. 2e) in conjunction with continuous MCx output during voluntary forelimb movements (a blue arrow in Fig. 2e) ^10^. Considering the fact that MCx output is transmitted to peripheral afferents (a red arrow in Fig. 2e), we hypothesized that MCx output might modulate muscles in a delayed manner via the transafferent pathway, which is composed of the descending and spinal reflex pathways (an orange arrow in Fig. 2f), in addition to direct control via the descending pathway (a blue arrow in Fig. 2f). Then, we sought to determine whether both the direct descending motor drive (descending input) and delayed action through the transafferent pathway from the MCx (transafferent input) contribute to the generation of muscle activity. We posited a delayed linear sum of descending and transafferent inputs as a first-order model of muscle activity. A reasonable conduction time for most peripheral afferent activity in DRGs to reach spinal motor neurons is 5-50 ms (see Methods). By simply adding the afferent conduction time (5-50 ms) to the time from MCx to afferents (70 ms) (Fig. 2e), we determined that most MCx activities require 75-120 ms to reach spinal motor neurons via the transafferent pathway (Fig. 2f). As mentioned, most MCx activity requires 5-50 ms to reach spinal motor neurons ^13, 17-19^. Therefore, we built a linear model to explain the instantaneous muscle activity using the descending input for 5-50 ms and the transafferent input for 75-120 ms preceding the timing of muscle activity to be calculated (Extended Data Fig. 2a). The model accurately reconstructed the overall temporal pattern of muscle activity, outperforming models constructed using shuffled controls (Extended Data Fig. 2b,c). Furthermore, the muscle activity calculated from both descending and transafferent inputs was reconstructed more accurately than that from the descending input alone (Extended Data Fig. 2d,e) or that from descending and shuffled transafferent inputs (Extended Data Fig. 2f,g). These results suggest that transafferent input is essential for the accurate reconstruction of muscle activity.

If MCx activity induces muscle activity via the transafferent pathway, which includes the spinal reflex pathway, the effects of the delayed action through the transafferent pathway from the MCx on muscles (an orange arrow in the top panel of Fig. 3a) must correspond to the effects of somatosensory feedback signals from peripheral afferents on muscles (a green arrow in the top panel of Fig. 3c). To investigate this, we decomposed the reconstructed muscle activity to identify the subcomponents of each input that affected the muscle activity. We calculated descending and transafferent components from the models built from the descending and transafferent inputs (Extended Data Fig. 3a,b). To assess the effects of somatosensory feedback signals on muscles, we similarly built a linear model in which the descending input and activity of peripheral afferents (afferent input) together accounted for the subsequent muscle activity and yielded the descending and afferent components from the models (Extended Data Fig. 3c,d). The temporal profile of the transafferent component was similar to that of the afferent component (Fig. 3a,c). To evaluate the similarity, we used shuffled data of the transafferent input and the MCx activity occurring 5–50 ms after the muscle activity to be calculated as controls for transafferent input and yielded the corresponding subcomponent (the shuffled and delayed components, respectively) (Extended Data Fig. 4). The similarity of the temporal profile (temporal similarity) between the afferent and transafferent components was higher than the temporal similarity between the afferent and shuffled components and between the afferent and delayed components (Fig. 3e). The temporal profile of the descending component in the model built from the descending and transafferent inputs was also similar to that of the descending component in the model built from the descending and afferent inputs (Fig. 3a,c,e).

**Fig. 3.**
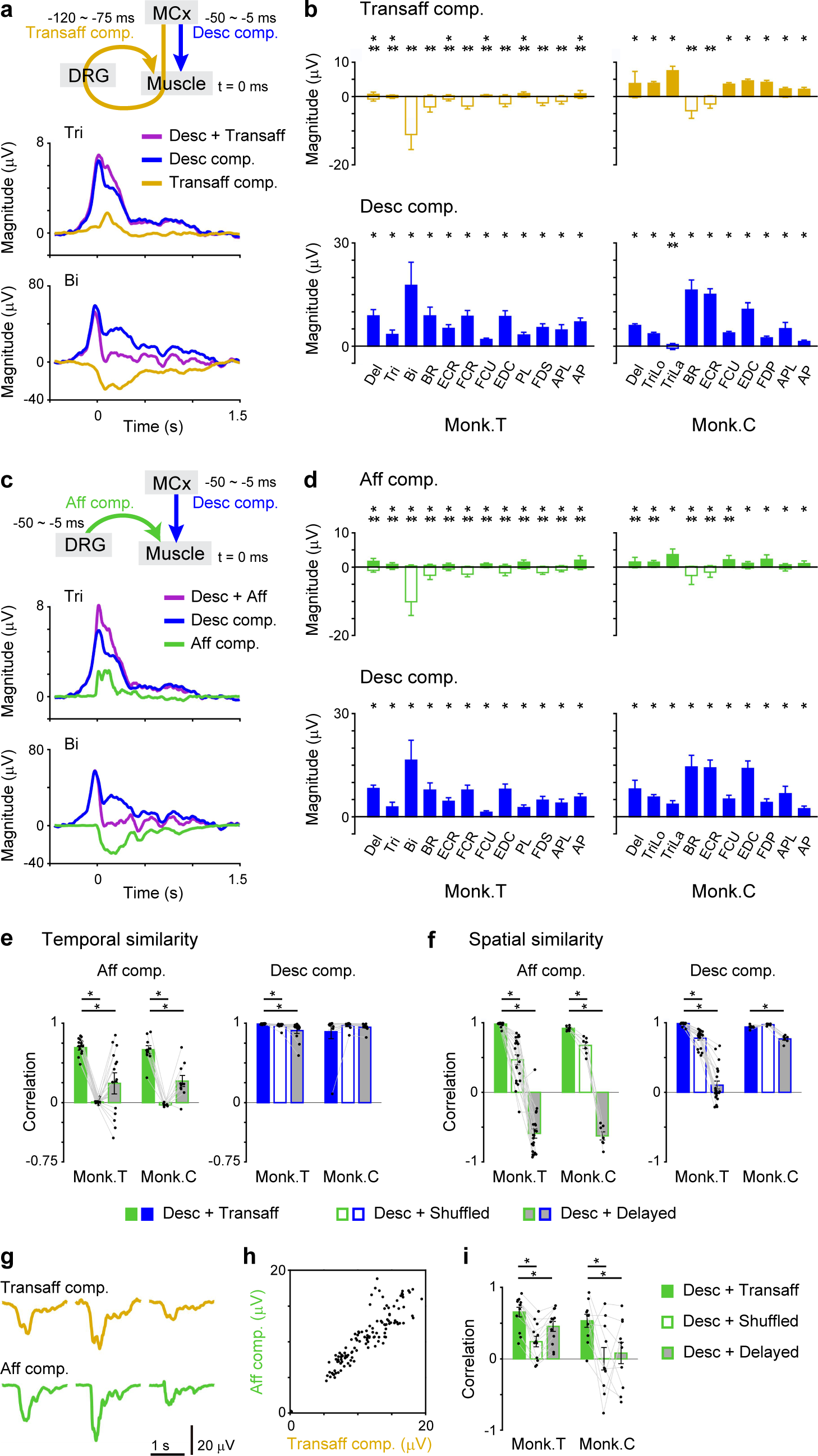
The MCx encodes the activation of muscles through the spinal reflex pathway. **a**, Reconstruction using descending and transafferent inputs and each subcomponent aligned to movement onset. **b**, Size of subcomponents (monkey T, 21 sessions; monkey C, 7 sessions). **c** and **d**, Same as in **a** and **b** for descending and afferent components, respectively. **e**, Temporal similarity between afferent and transafferent components (left) or across descending components (right) for different models (monkey T, 12 muscles; monkey C, 10 muscles). **f**, Same as in **e** for the spatial similarity of afferent (left) or descending (right) components across muscles for different models (monkey T, 21 sessions; monkey C, 7 sessions). **g**, Transafferent and afferent components in three datasets. **h**, Scatter plots of the size of transafferent versus afferent components. Dot, a single dataset. **i**, Average correlation between afferent and transafferent components in the scatter plot for the different models (monkey T, 12 muscles; monkey C, 10 muscles). In **b** and **d**, data are the mean ± SD. * and \*\**P* < 0.05, unpaired two-tailed *t*-test for positive and negative values. In **e**, **f**, and **i**, data are the mean ± SEM. *P* < 0.05, one-way repeated-measures analysis of variance [ANOVA]; \**P* < 0.05, paired two-tailed *t*-test.

The size of the subcomponents varied across different muscles. The distribution of the transafferent component size across different muscles was similar to that of the afferent component size (Fig. 3b,d). The similarity of the distribution of afferent and transafferent components across muscles (spatial similarity) was higher than the spatial similarity between the afferent and shuffled components and between the afferent and delayed components (Fig. 3f). The spatial profile of the descending component in the model built from the descending and transafferent inputs was also similar to that of the descending component in the model built from the descending and afferent inputs (Fig. 3b,d,f). These results suggest that MCx activity between −75 and −120 ms before muscle activity encoded similar spatiotemporal information about muscle activity as the subsequent afferent activity.

The size of the afferent component was variable across datasets (Fig. 3g). If the effects of transafferent input on muscles correspond to afferent input, the variation in the size of the transafferent component across datasets should reflect the variation in the size of the afferent component. We found a positive correlation between the size of the afferent and transafferent components (Fig. 3h). The correlation between the size variation of the afferent and transafferent components was notably larger than those between the afferent and shuffled components and between the afferent and delayed components (Fig. 3i). Taken together, these findings suggest that MCx activity from −120 to −75 ms and from −50 to −5 ms before muscle activity encoded the muscle activity evoked by the afferent and descending inputs, respectively.

## MCx encodes the spinal reflex

Next, we wondered whether activation of limb muscles through the transafferent pathway corresponds to the action of spinal reflexes on limb muscles, such as the stretch reflex and reciprocal inhibition. Our previous study analyzing the relationship between initial limb movement and subsequent afferent components indicated that afferent effects on muscle activity conform to the pattern of spinal reflexes ^10^. For instance, monkey C initiated reaching by supination of the elbow joint and flexion of the wrist joint (Fig. 4a). The afferent component at the beginning of the reaching movement (55 to 100 ms around movement onset) exhibited a facilitatory effect on the elbow extensor, the triceps brachii lateralis (TriLa), and a suppressive effect on the elbow flexor, the brachioradialis (BR), which act as an antagonist of the TriLa (Fig. 4b,c). These antagonistic activations on extensor and flexor muscles could be attributed to the stretch reflex and reciprocal inhibition, respectively. The results obtained from monkey T also indicated a similar relationship between initial joint movements and corresponding afferent components (Fig. 4c). Then, we investigated whether the transafferent effects on muscle activity can also be attributed to the stretch reflex and reciprocal inhibition. We calculated the transafferent component at the beginning of the reaching movement. Similar to afferent components, transafferent input exerted a facilitatory effect on some agonist muscles (elbow extensors) and a suppressive effect on antagonist muscles (elbow and wrist flexors in monkey T and an elbow flexor in monkey C) (Fig. 4c). We calculated the shuffled or delayed components during the same period, but the results were not consistent with spinal reflex action (Extended Data Fig. 5). Therefore, the effects of transafferent inputs on muscle activity are at least partially accounted for by the action of spinal reflexes. These results suggest that the MCx encodes the spinal reflex.

**Fig. 4.**
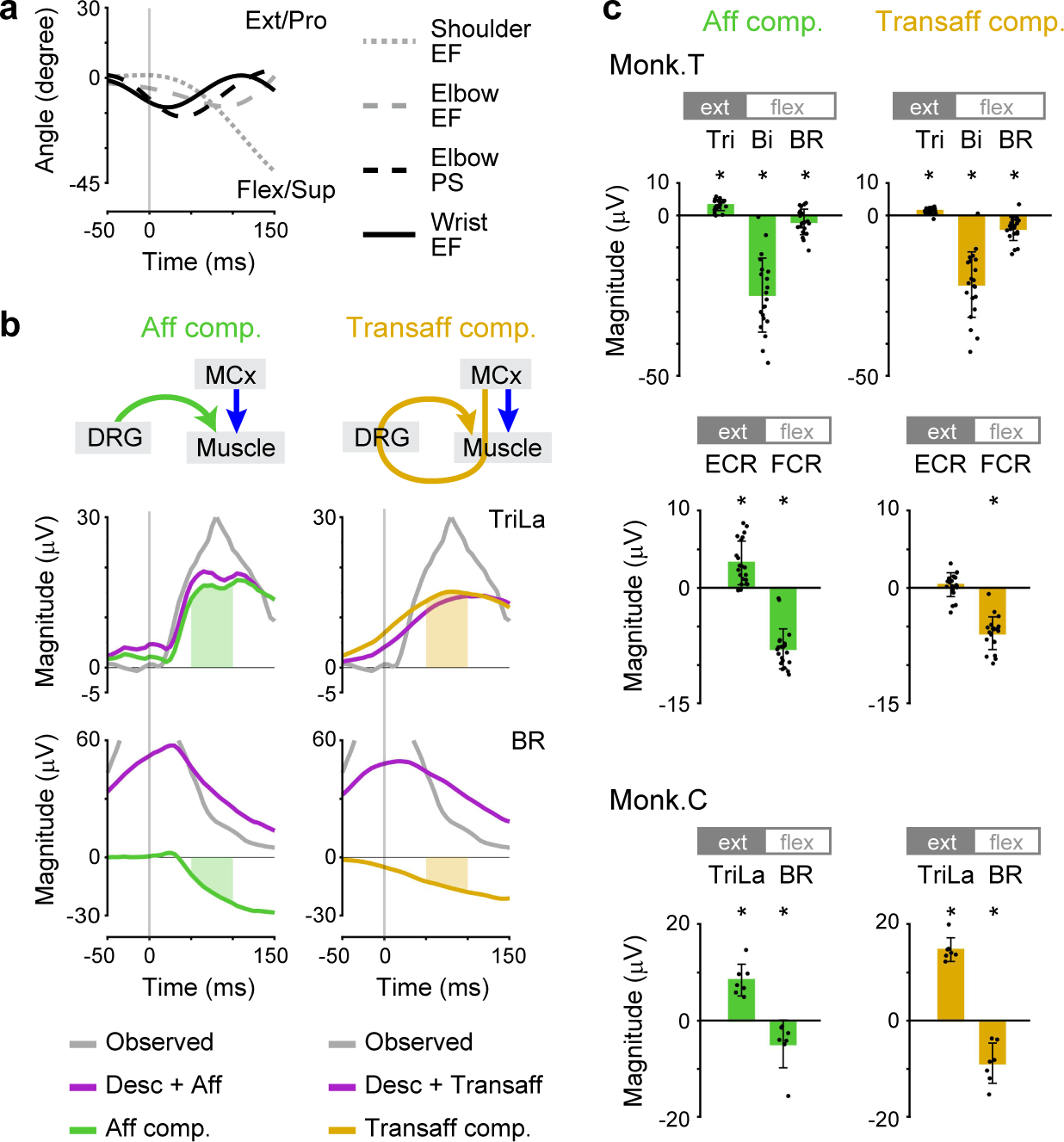
The effects of transafferent input on muscles are accounted for by the stretch reflex and reciprocal inhibition. **a**, Forelimb joint angles of monkey C when the monkey began to reach. **b**, *Left*: The observed muscle activity of monkey C, its reconstruction using descending and afferent inputs, and the afferent component of the reconstruction. The vertical lines indicate the time of movement onset. *Right*: The observed muscle activity, its reconstruction using descending and transafferent inputs, and transafferent component in the reconstruction. **c**, Size of afferent (left column) and transafferent (right column) components for antagonistic muscle pairs (ext: extensor; flex: flexor) in a period from the beginning of the reaching movement (55 to 100 ms around movement onset; shown in the green and orange areas in **b**, monkey T, 21 sessions; monkey C, 7 sessions). In **c**, data are the mean ± SD. \**P* < 0.05, unpaired two-tailed *t-*test.

## M1 is the major source of the spinal reflex

The corticospinal projections of primates play an important role in relaying motor commands from multiple motor-related areas to the spinal cord ^2^. Next, we asked which cortical area contributes to the generation of muscle activity through the transafferent pathway. By recording ECoG signals in the PMd, PMv, and M1 with a multichannel electrode array, we were able to compare the effective activity across these areas. We calculated the descending and transafferent components based on the activity in each cortical area and found that the subcomponents calculated from M1 activity were much more prominent than the subcomponents calculated from PMd or PMv activity (Fig. 5a-c and Extended Data Fig. 6a). Similar to this result, the subcomponents calculated from M1 activity for the reconstruction of afferent activity were also more prominent than the subcomponents calculated from PMd or PMv activity (Fig. 5a-c and Extended Data Fig. 6a). M1 sites with the largest subcomponent for the transafferent input, which corresponds to the action of spinal reflexes, were located just anterior to the central sulcus, as was the case for the descending input and the afferent reconstruction (Fig. 5d and Extended Data Fig. 6b). Thus, the transafferent input from a subset of M1 rather than the PMd and PMv could primarily account for muscle activity.

**Fig. 5.**
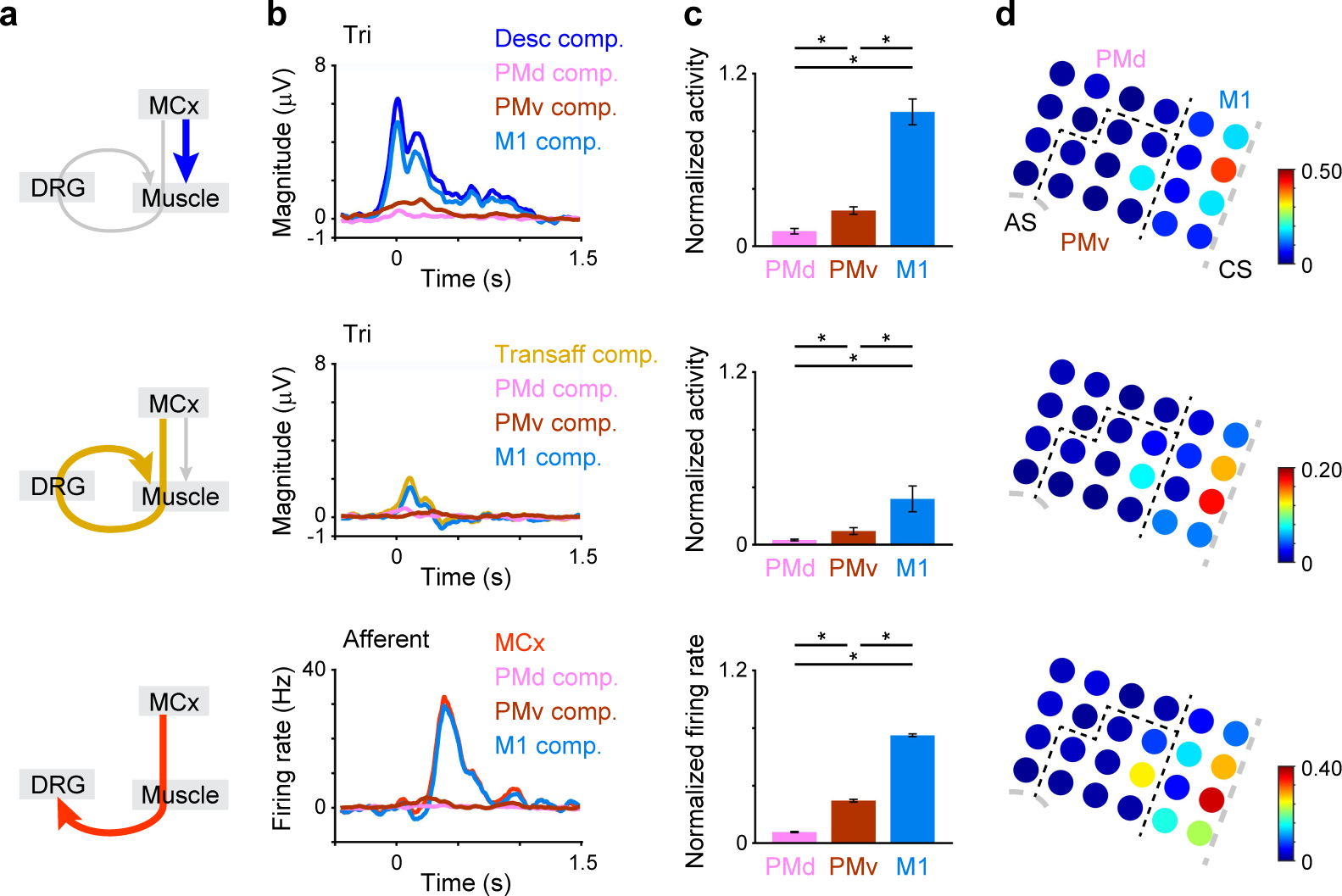
M1 is a major predictor of muscle activity by descending and transafferent pathways. **a**, The thick, colored arrows show the inputs for which the modulation is represented in **b-d**. **b**, *Top and Second*: Reconstruction of muscle activity using descending and transafferent inputs and subcomponents calculated from the activity in each cortical area aligned to the movement onset of monkey T. *Bottom*: Reconstruction of afferent activity using MCx activity and subcomponents calculated from the activity in each cortical area aligned to the movement onset of monkey T. **c**, Size of subcomponents calculated from the activity in each cortical area for the prediction of muscle and afferent activity (monkey T, 12 muscles or 702 afferents). **d**, Color maps represent the size of subcomponents calculated from the activity at each electrode for the prediction of muscle or afferent activity in monkey T. The size of each subcomponent is normalized by the size of the reconstruction of muscle activity using the descending and transafferent inputs or the reconstruction of afferent activity using the descending input. CS, central sulcus; AS, arcuate sulcus. In **c**, data are the mean ± SEM. *P* < 0.05, one-way repeated-measures ANOVA, \**P* < 0.05, paired two-tailed *t*-test.

## Transafferent M1 output drives muscles

If M1 signals transmitted by the transafferent pathway are crucial for driving muscle activity, selective blockade of the transafferent pathway should result in a reduction in muscle activity, especially its transafferent component. We tested this possibility by sectioning peripheral afferents (i.e., performing dorsal rhizotomy) at the lower cervical level (C6-C8), which innervate the forearm, in two other monkeys. Dorsal rhizotomy causes the large-scale reorganization of neural circuits through axonal sprouting, which takes several weeks ^20, 21^. To minimize such long-term effects on neural reorganization, we analyzed the data obtained immediately after dorsal rhizotomy (up to 5 days after dorsal rhizotomy). Attenuation of somatosensory evoked potentials (SEPs) elicited by electrical stimulation of forelimb muscles indicated a blockade of most signal transmission from peripheral afferents (Fig. 6a). Although the monkeys showed loss of dexterous finger movements after rhizotomy, they were able to reach the target. We recorded activity in the MCx and the forelimb muscles when the monkeys performed reaching movements before and after dorsal rhizotomy (Extended Data Fig. 7). The total muscle activity and the peak amplitude of muscle activity during movements decreased after dorsal rhizotomy (gray lines in Fig. 6b-d and Extended Data Fig. 8a). This result suggested that afferent inputs transmitted through the somatic reflex arc contribute to the activation of forelimb muscles during voluntary movements.

**Fig. 6.**
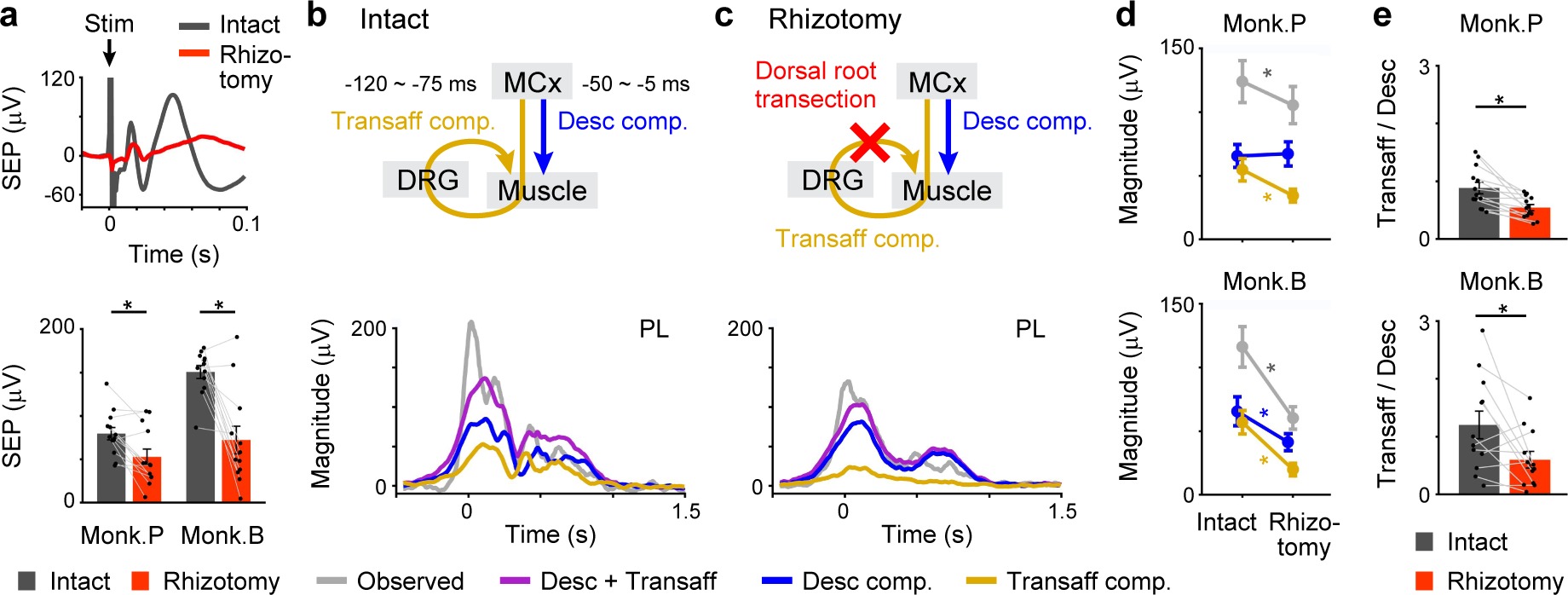
Sectioning of peripheral afferents impairs the transafferent activation of muscles by the MCx. **a**, *Top*: Example of somatosensory evoked potentials (SEPs) elicited by electrical stimulation of a forelimb muscle (EDC). *Bottom*: Sizes of SEPs recorded over the primary somatosensory cortex (monkey P, 13 muscles; monkey B, 12 muscles). **b**, The observed muscle activity, its reconstruction using descending and transafferent inputs, and each subcomponent aligned to movement onset before dorsal rhizotomy. **c**, Same as in **b**, but after dorsal rhizotomy. **d**, Total muscle activity (gray) and size of descending (blue) and transafferent (orange) components before and after dorsal rhizotomy. The muscle activity and the size of descending and transafferent components after dorsal rhizotomy were compared to those observed before dorsal rhizotomy (monkey P, 13 muscles; monkey B, 12 muscles). **e**, Ratio of the size of the transafferent component to that of the descending component was compared before and after dorsal rhizotomy (monkey P, 13 muscles; monkey B, 12 muscles). In **a**, **d**, and **e**, data are the mean ± SEM. \**P* < 0.05, paired two-tailed *t*-test.

Next, to examine whether dorsal rhizotomy affects the transafferent descending activation of muscles, we built a linear model using descending and transafferent inputs to account for the subsequent muscle activity and calculated the descending and transafferent components. The model reconstructed the overall temporal pattern of muscle activity before and after dorsal rhizotomy more accurately than the shuffled control (Extended Data Fig. 8b,c). Among the motor-related areas, M1 encodes most information on muscle activity transmitted via the direct descending and transafferent pathways in monkeys P and B (Extended Data Fig. 9). We calculated the size of the transafferent component during movement and found that it decreased after dorsal rhizotomy (Fig. 6b-d). This result demonstrated that dorsal rhizotomy impaired signal transmission from the M1 through the transafferent pathway, which usually activates muscles in intact animals. On the other hand, the size of the descending component remained after dorsal rhizotomy in monkey P (Fig. 6d). Furthermore, the ratio of the size of the transafferent component to that of the descending component decreased after dorsal rhizotomy in monkeys P and B (Fig. 6e), indicating that the decrease in the transafferent component size cannot be explained by the decrease in the descending component size alone. These results provide causal evidence that M1 signals transmitted through the transafferent pathway are involved in the activation of forelimb muscles during voluntary movements.

## Discussion

How supraspinal and spinal structures in the nested hierarchy control limb muscles is a pivotal question in the neural control of limb movements. The present study demonstrates that M1 drives muscle activity not only through direct activation via the descending pathway but also through delayed activation via the transafferent pathway (Extended Data Fig. 10). Thus, to ensure accurate control of limb muscles through two pathways that operate on different time scales, M1 provides appropriate motor output based on an internal forward model that prospectively predicts future spinal reflexes.

Multiple pathways exist for signals to travel from the M1 to peripheral afferents. The most relevant time difference between the M1 and afferent activity is 70 ms (Fig. 2d). Based on the assumption that afferent activity requires approximately 50 ms to be induced by the activation of limb muscles ^22^, it is likely that activation of alpha motor neurons and ensuing limb movements lead to afferent activity, including signals from cutaneous receptors in skin and proprioceptors in muscles, tendons, and joints. Muscle contraction also triggers Golgi tendon organ activation. In addition, afferent activity was detected around the timing of movement onset (Extended Data Fig. 1), which may result from the gamma efferent drive leading to muscle spindle activation. The strong relationship between the M1 and peripheral afferents of various modalities could potentially underlie the encoding of the transafferent activation of muscles by the M1.

Direct corticospinal projections from the M1 in primates are the densest and most numerous among motor-related cortical areas ^23^. Additionally, the most notable feature in the primate M1 is the presence of direct connections from pyramidal neurons in the M1 to spinal motor neurons, which exert a strong impact on muscle activity ^24-29^. It is therefore widely assumed that these robust descending control signals from the M1 act on spinal motor neurons to produce the appropriate muscle activity required to execute the intended movement ^15^. Our study demonstrated that M1 primarily encoded muscle activity driven by transafferent pathways during movements (Fig. 5 and Extended Data Figs. 6 and 9), which was decreased after sectioning dorsal roots at the cervical segments (Fig. 6b-e). These findings suggest that the impact of M1 on spinal motor neurons further activates spinal reflexes.

Dorsal rhizotomy reduced muscle activity and the transafferent component (Fig. 6b-e and Extended Data Fig. 8a). These findings suggest that spinal reflexes amplify the direct descending commands from the M1 in the intact condition. Hence, the M1 minimizes cortical energy expenditure by leveraging the spinal reflex system to control limb muscles adequately. This efficient control of limb muscles by M1 is consistent with the theory of optimal feedback control, a leading theoretical framework in motor control, which computes the optimal strategy for reducing movement error and motor effort ^30, 31^. According to this theory, the control policy provides feedforward motor commands to the spinal motor neurons based on the estimated current state and task demands. The MCx, brainstem, and peripheral afferents projecting to spinal motor neurons are assumed to be the brain regions responsible for the control policy ^32^. How the distributed organization of the control policy controls the end effectors is not yet fully understood ^33^. The current study showed that the M1 prospectively computes sensorimotor integration in the spinal cord in voluntary limb movement, providing evidence for the neuronal mechanism underlying the interplay between the M1 and peripheral afferents. Incorporating spinal model implementation by the M1 into a theory for movement control will help us to clarify how the central nervous system controls voluntary limb movements.

## Acknowledgments

We thank Y. Yamanishi for animal care, Y. Nishihara and M. Togawa for technical help, Professor K. Seki for encouragement, and Professor A. Pruszynski and Professor S. Perlmutter for helpful discussions and critical comments. The monkeys used in this study were provided by the National Bioresource Project of the Ministry of Education, Culture, Sports, Science, and Technology of Japan (MEXT). This work was performed with support from grants from the Medtronic Japan External Research Institute, the Takeda Science Foundation and the Japan Agency for Medical Research and Development to T.U., a Grant-in-Aid for Scientific Research to T.I. (19H01011, 19H05723), and the Strategic Research Program for Brain Sciences from MEXT, the Japan Agency for Medical Research and Development, the Japan Science and Technology Agency, a Grant-in-Aid for Scientific Research from MEXT to Y.N. (23680061, 25135733).

## Author contributions

Design of the experiments related to simultaneous recordings: TU, YN. Conducting the experiments related to simultaneous recordings: TU, YN. Analyzing the data related to simultaneous recordings: TU. Design of the experiments related to dorsal rhizotomy: OY, TU, YN. Conducting the experiments related to dorsal rhizotomy: OY, MS, MK, YN. Analyzing the data related to dorsal rhizotomy: TU, OY. Interpreting the data: TU, OY, TI. YN. Writing – original draft: TU, OY, TI. YN. Writing – review & editing: TU, OY, MS, MK, TI, YN.

## Competing interests

The authors declare no competing interests.

## Data and materials availability

All data needed to evaluate the conclusions presented in this manuscript are included in the manuscript and/or in the Extended Data section. Additional data related to this manuscript may be requested from the authors.

## Methods

### Animals

We used one adult male monkey (monkey T, weight 6–7 kg, *Macaca fuscata*) and three adult female monkeys (monkey C, weight 5–6 kg, *Macaca mulatta*; monkey P, weight 4*–*5 kg, *Macaca fuscata*; and monkey B, weight 5–6 kg, *Macaca fuscata*). The experiments were approved by the Experimental Animal Committee of the National Institute of Natural Sciences and Tokyo Metropolitan Institute of Medical Science. The animals were cared for and treated humanely in accordance with National Institutes of Health guidelines. Part of the dataset obtained from monkeys T and C is the same as the dataset used in our previous studies ^10, 13^.

### Surgery

All surgical procedures were performed using sterile techniques while the animal was anesthetized with 1–2% isoflurane (monkeys T and C) or sevoflurane (monkeys P and B). Dexamethasone, atropine, and ampicillin were administered preoperatively; ampicillin and ketoprofen were given postoperatively.

For EMG recordings, we implanted pairs of Teflon-insulated wire electrodes (AS631; Cooner Wire) into the forelimb muscles on the right side. We used the activity in the deltoideus posterior (Del), triceps brachii (Tri), biceps brachii (Bi), brachioradialis (BR), extensor carpi radialis (ECR), flexor carpi radialis (FCR), flexor carpi ulnaris (FCU), extensor digitorum communis (EDC), palmaris longus (PL), flexor digitorum superficialis (FDS), abductor pollicis longus (APL), and adductor pollicis (AP) of monkey T; the Del, triceps brachii longus (TriLo), TriLa, BR, ECR, FCU, EDC, flexor digitorum profundus (FDP), APL, and AP of monkey C; the Del, Tri, Bi, BR, ECR, ECU (extensor carpi ulnaris), FCU, EDC, ED23 (extensor digitorum-2,3), ED45 (extensor digitorum-4,5), PL, FDS, and FDP of monkey P; and the Del, Tri, Bi, BR, ECR, FCR, ECU, EDC, ED23, PL, FDS, and AP of monkey B.

To record ECoG signals from the MCx, we implanted a 32-channel (monkeys T and C) or 30-channel (monkeys P and B) grid electrode array (Unique Medical) with a diameter of 1 mm and an interelectrode distance of 3 mm beneath the dura mater over the sensorimotor cortex (Extended Data Figs. 6b and 9b). We placed the ground and reference electrodes over the ECoG electrode so that they contacted the dura.

To record afferent signals, we implanted two multielectrode arrays (Blackrock Neurotech) into the dorsal root ganglia at the cervical level (monkey T, C7 and C8; monkey C, C6 and C7) on the right side. To block peripheral afferent inputs, we sectioned the cervical dorsal rootlets (dorsal rhizotomy) at segments C6-C8 ^34, 35^.

### Behavioral task

All monkeys were operantly conditioned to perform a reach-to-grasp task with the right hand (Fig. 1a). After putting its hand on a home button for 2–2.5 s (monkeys T and C) or 1 s (monkeys P and B), the monkey reached for a lever and pulled it to receive a reward.

### Recordings

All neural and muscular signals were recorded simultaneously using a data acquisition system (Plexon for monkeys T and C and Tucker-Davis Technologies for monkeys P and B).

EMG signals were amplified using amplifiers (AB-611J; Nihon Kohden); they were sampled at 2,000 Hz in monkey T and at 1,000 Hz in monkey C at a gain of ×1,000–2,000 and sampled at 1,017.3 Hz in monkeys P and B at a gain of ×100. We applied a 2nd-order Butterworth bandpass filter (1.5–60 Hz) to the signals, rectified the filtered signals, resampled the signals to 200 Hz, and smoothed the resampled signals using a moving window of 11 bins.

The ECoG signals of monkeys T and C were amplified using a multichannel amplifier (Plexon MAP system; Plexon) at a gain of ×1,000 and sampled at 2,000 Hz in monkey T and at 1,000 Hz in monkey C. The ECoG signals of monkeys P and B were amplified using a multichannel amplifier (PZ2; Tucker-Davis Technologies) at a gain of ×51 and sampled at 1,017.3 Hz. We applied a 2nd-order Butterworth bandpass filter (1.5–240 Hz) to the signals, computed a short-time fast Fourier transform on moving 100-ms windows of the filtered signals at a 5-ms step, normalized the power to the average power in each session, and calculated the average power in the high-gamma bands (high-gamma 1, 60–120 Hz; high-gamma 2, 120–180 Hz). We considered the high-gamma power of the ECoG signals to be representative of neural activity in cortical areas ^14^.

The peripheral afferent activities of monkeys T and C were initially amplified at a gain of ×20,000 and sampled at 40 kHz (Plexon MAP system; Plexon). We extracted filtered waves (150–8,000 Hz) above an amplitude threshold, sorted the thresholded waves using semiautomatic sorting methods (Offline Sorter; Plexon), and performed manual verification. We isolated 25–39 units in monkey T and 11–15 units in monkey C. We convolved the inversion of the interspike interval using an exponential decay function whose time constant was 50 ms and resampled the firing rate to 200 Hz.

We calculated the movement-related modulation of the EMG signals, the ECoG signals, and the peripheral afferent activity before analyzing the data. We first calculated the baseline activity by averaging the activity from −1,250 to −750 ms around movement onset. We then subtracted the baseline activity from the preprocessed activity. We used movement-related modulation throughout the premovement and movement periods (−500 to 1,500 ms around movement onset) for monkeys T, C, and P as a single trial for further analysis. Since monkey B could reach the lever but did not grasp it after dorsal rhizotomy, we used movement-related modulation throughout the premovement and movement periods (−500 to 250 ms around movement onset) for monkey B.

We recorded the times at which the animals released the home button, pulled the lever, and pushed the home button.

We recorded the forelimb movements of monkeys T and C using an optical motion capture system with 12 cameras (Eagle-4, Motion Analysis). The spatial positions of ten reflective markers attached to the surface of the forelimbs and body were sampled at 200 Hz.

We assessed the sectioning of peripheral afferents by recording SEPs over the primary somatosensory cortex elicited by electrical stimulation (2 mA, 200 monophasic pulses at 2.42 Hz) of the forelimb muscles. To calculate SEP size, we subtracted the minimum value that was recorded between 11 ms and 20 ms after stimulation by any of the electrodes over the primary somatosensory cortex from the maximum value that was recorded between 16 ms and 25 ms after stimulation.

### Sparse linear regression

Neural ensemble activity in M1 satisfactorily accounts for muscle activity when a linear model is used ^36, 37^. We examined whether the integration of the descending signals from M1 and somatosensory signals from peripheral afferents in spinal motor neurons can also be represented as a linear relationship. We modeled muscle activity as a weighted linear combination of high-gamma activity in the MCx and/or neuronal activity of peripheral afferents using multidimensional linear regression as follows:

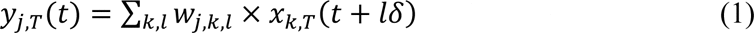

where *y_j,T_(t)* is a vector of the EMG activity of muscle *j* (12, 10, 13, and 12 muscles for monkeys T, C, P, and B, respectively) at time index *t* in trial *T*, *x_k,T_(t+lδ)* is an input vector of the peripheral afferent or cortical signal *k* at time index *t* and lag time *lδ* (*δ* = 5 ms, *l* = −10 – −1) in trial *T*, and *w_j,k,l_* is a vector of weights on the peripheral afferent or cortical signal *k* at lag time *lδ*. We applied a Bayesian sparse linear regression algorithm that introduces sparse conditions for the unit/channel dimension ^38^. As we examined how the combined activity in the MCx and/or peripheral afferents influenced subsequent muscle activity, lag time *lδ* (Equation 1) was set to negative values. To represent the direct effect of the MCx on muscle activity through the descending pathway (descending input), we used activity in the MCx from −50 to −5 ms to reconstruct muscle activity at time 0 for the following reasons. Averaging the muscle activity triggered at the spiking activity of M1 neurons shows postspike facilitation with a latency of 6.7 ms ^24^. Accordingly, we set 5 ms as the shortest lag time. The weighted sum of M1 neuronal activity accurately accounted for muscle activity at a lag time of 40–60 ms ^39^. However, it is possible that the MCx has some effect on muscle activity through the somatic reflex arc within 70 ms, as shown in Figure 2d. To avoid the influence of the somatic reflex arc, we set 50 ms as the longest lag time. To represent the effect of peripheral afferents on muscle activity (afferent input), we used the activity in the peripheral afferents from −50 to −5 ms to reconstruct muscle activity at time 0 for the following reasons. Averaging the muscle activity triggered at the spiking activity of peripheral afferents showed postspike facilitation with a latency of 5.8 ms ^40^. Thus, we set 5 ms as the shortest lag time. In addition, the delay time in the reflex pathways was reported to be 47 ms ^41^. The transcortical long-latency reflex is detected as a burst of muscle activity occurring 50–100 ms following an imposed limb displacement ^42^. To avoid the influence of the transcortical long-latency reflex on muscle activity, we set 50 ms as the longest lag time. To represent the reafferent effect of the MCx on muscle activity through the transafferent pathway composed of descending and spinal reflex pathways (transafferent input), we used the activity in the MCx from −75 to −120 ms to reconstruct muscle activity at time 0. We obtained this lag time by simply adding the time at which the MCx most accurately accounted for the peripheral afferent activity (70 ms) to the lag time of the descending input.

To compute the contribution of each descending, afferent, and transafferent input to the reconstruction of muscle activity, we calculated each subcomponent of the reconstructed activity using the relevant input and the respective weight values in a decoding model built from the combined inputs. For example, the descending component was calculated as follows:

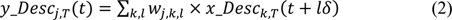

where *y_Desc_j,T_*(*t*) is a vector of the descending component at muscle *j* at time index *t* in trial *T*, *x_Desc_k,T_*(*t+lδ*) is an input vector of the cortical signal *k* at time index *t* and lag time *lδ* in trial *T*, and *w_j,k,l_* is derived from a vector of weights in Equation (1) but with the weights assigned to peripheral afferents removed. We also calculated subcomponents from the activity in each cortical area or each electrode in a similar way.

The activity of peripheral afferents was modeled as a weighted linear combination of MCx activity using multidimensional linear regression as follows:

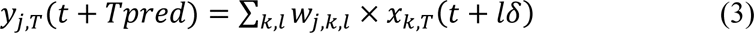

where *y_j,T_(t)* is a vector of the peripheral afferent *j* at time index *t* in trial *T*, *x_k,T_(t+lδ)* is an input vector of the cortical signal *k* at time index *t* and lag time *lδ* (*δ* = 5 ms, *l* = −10 to −1) in trial *T*, and *w_j,k,l_* is a vector of weights on the cortical signal *k* at lag time *lδ*. We considered that MCx activity evokes muscle activity that then generates peripheral afferent activity. We set the lag time *lδ* to negative values. By changing *Tpred* from 0 ms to 120 ms, we identified *Tpred*, the time point at which the reconstruction accuracy of the activity in peripheral afferents was most significant (70.7 ms, with an average of 62.7 ms for monkey T and 78.6 ms for monkey C, Fig. 2d).

### Data analysis

We analyzed data from 21 and 7 sessions, each of which was 10 min, for monkeys T and C, respectively. The movement times of monkeys P and B increased temporarily after dorsal rhizotomy. To compare the magnitude of muscle activity without the bias of differences in movement time, we used trials in which the monkeys performed the movement within a certain period (0.65–0.85 s for the movement time of monkey P and 0.13–0.19 s for the reach time of monkey B) based on the distribution of movement or of reach time observed for these animals in the intact condition. We divided the data into datasets that included no fewer than 129 trials (see below). For monkey P, we analyzed data from 17 and 4 sessions before and after dorsal rhizotomy, respectively; for monkey B, we analyzed data from 8 and 2 sessions before and after dorsal rhizotomy, respectively. We used data obtained up to 5 days after dorsal rhizotomy to examine the acute effect of dorsal rhizotomy without the reorganization of neural circuits.

We built models designed to reconstruct the temporal changes in the EMG signals or afferent activity using a partial dataset (training dataset) and tested them using the remainder of the same dataset (test dataset). One hundred and eight trials were randomly selected as a training dataset, and 21 trials were selected randomly from the remaining trials as the test dataset. To assess the model, we calculated the correlation coefficients between the observed data and their reconstruction in the test dataset. We also calculated the variance accounted for (VAF) as follows:

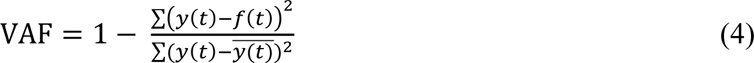

where *y*(*t*) is a vector of the actual activity in muscles at time index *t*, 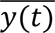 is the mean of *y(t)*, and *f(t)* is the reconstructed activity at time index *t*. We performed 6-fold cross-validation in the analysis of each session and used averaged values for the analysis. We then calculated the average activity of each muscle or peripheral afferent using data taken from 21 (monkey T), 7 (monkey C), 17 (monkey P, before rhizotomy), 4 (monkey P, after rhizotomy), 8 (monkey B, before rhizotomy), or 2 (monkey B, after rhizotomy) sessions. In control analyses of the model reconstruction, we created surrogate training datasets in which we shuffled the temporal profiles of the inputs independently across different blocks to generate a model, and we subsequently calculated the reconstruction using unshuffled data to evaluate the model (Fig. 2b,c and Extended Data Figs. 2b,c and 8b,c) or shuffled data to evaluate the inputs (Fig. 3e,f,i and Extended Data Figs. 4c,d and 5). We also used the MCx activity occurring 5–50 ms after the muscle activity to be calculated as another control dataset for evaluation of transafferent inputs (Fig. 3e,f,i and Extended Data Figs. 4e,f and 5).

We observed a period in which there was movement-related modulation of muscles. To obtain the onset time of the observed muscle activity, we first calculated the average of the aligned waveforms in the test dataset. We then defined one-fifth of the maximum amplitude of the observed muscle activity from 250 ms before to 250 ms after movement onset as a threshold. If the activity or the reconstruction exceeded the threshold in 5 consecutive bins, the first of these bins was set as the onset. We calculated the average onset values observed in six test datasets in one session and obtained their average values over all sessions. To obtain offset time, we calculated the average of the aligned waveforms of the muscle activity and analyzed data obtained 500 ms after the onset of movement. If the averaged data values were below the same threshold that was used to calculate the onset time in 5 consecutive bins, the first of these bins was set as the offset. The calculated onset and offset corresponded well with those determined by visual inspection. We calculated the area above or below baseline for each component during the period in which movement-related modulation of muscles was detected (monkey T, −100 to 1,150 ms around movement onset; monkey C, −100 to 1,300 ms; and monkey P, −200 to 800 ms) and from the time of onset of muscle activity to the time the animal reached the lever (monkey B, −300 to 150 ms) (Figs. 3b,d and 6d,e and Extended Data Figs. 4b,d,f). We similarly calculated the sum of areas above and below baseline for each component from each cortical area and each electrode during these periods and normalized them to the values from the whole area (Fig. 5c,d and Extended Data Figs. 6 and 9). Then, we divided these areas by the time of movement-related modulation of muscles. We calculated the firing rate of afferents and that of each component from each cortical area and each electrode during the period in which movement-related modulation of muscles was detected (monkey T, −100 to 1,150 ms around movement onset; monkey C, −100 to 1,300 ms) and divided the values by the time of movement-related modulation of muscles (Fig. 5c,d and Extended Data Fig. 6). We tested whether the positive or negative values deviated from zero using a standard *t*-test. The time at which the wrist joint angle initially peaked along the flexion/extension direction of monkey T was 51.4 ± 9.6 ms (mean ± SD), and the times at which the wrist joint angle initially peaked along the flexion/extension direction and at which the elbow joint angle initially peaked along the pronation/supination direction of monkey C were 31.4 ± 5.6 ms and 47.1 ± 2.7 ms, respectively (Fig. 4a). We calculated the temporal mean of each component during the period from the beginning of the reaching movement (from 55 to 100 ms around movement onset) (Fig. 4c and Extended Data Fig. 5). We statistically tested whether the temporal mean deviated from zero using a standard *t*-test.

We obtained the lag time between afferents and MCx activity at which the afferent activity was most accurately reconstructed from MCx activity using a linear model. We first selected data for which the correlation coefficient or VAF of the model was superior to that of models built using a surrogate shuffled control. We then created a graph of the relationship between the lag time and reconstruction accuracy using only the data that led to high accuracy (correlation coefficient greater than 0.4, VAF greater than 0.15). We fitted a quadratic function to the graph and obtained the vertex of the fitting curve.

### Statistical analysis

We used the paired or unpaired Student’s *t*-test. When comparing more than 2 group means, we first assessed the data using one*-*way analysis of variance (ANOVA). The alpha level of significance was set at 0.05. The results of all statistical tests (including *P* values) are included in the Extended Data Tables. The data are expressed as the mean ± the standard error of the mean (SEM) or the mean ± SD. We used MATLAB R2020a (MathWorks) for statistical analysis. No statistical methods were used to predetermine the sample size. However, sample sizes followed published standards.

---

**Extended Data Fig. 1.**
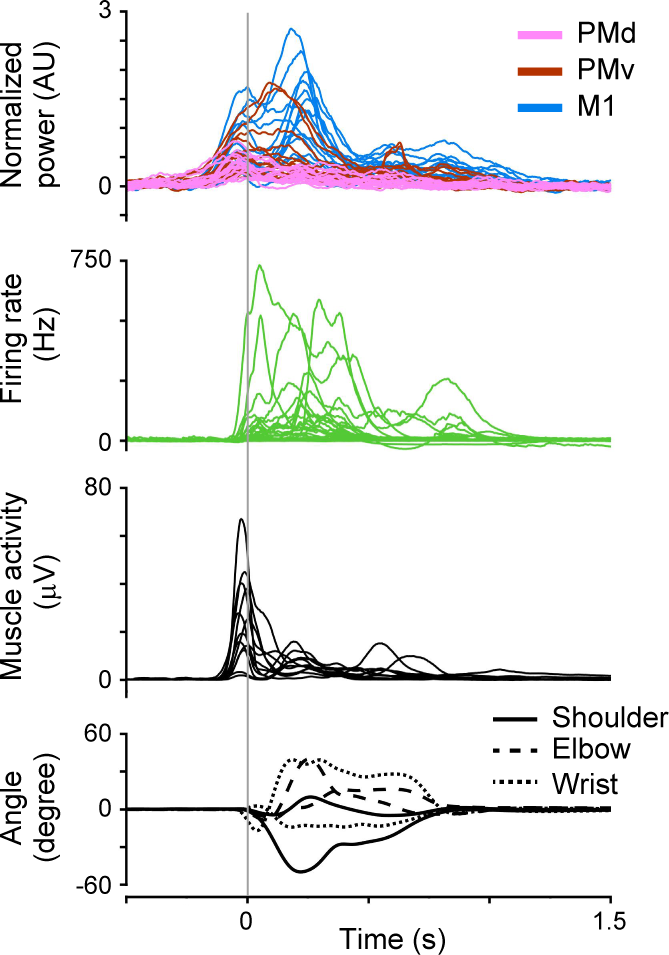
Simultaneous recording of MCx signals, afferent and muscle activity, and joint kinematics. Modulation of cortical and peripheral activity in monkey T aligned to movement onset. *Top*: High-gamma cortical activity. *Second*: Instantaneous firing rate of peripheral afferents. *Third*: Forelimb muscles. *Bottom*: Joint angles. A vertical line represents the time of movement onset.

**Extended Data Fig. 2.**
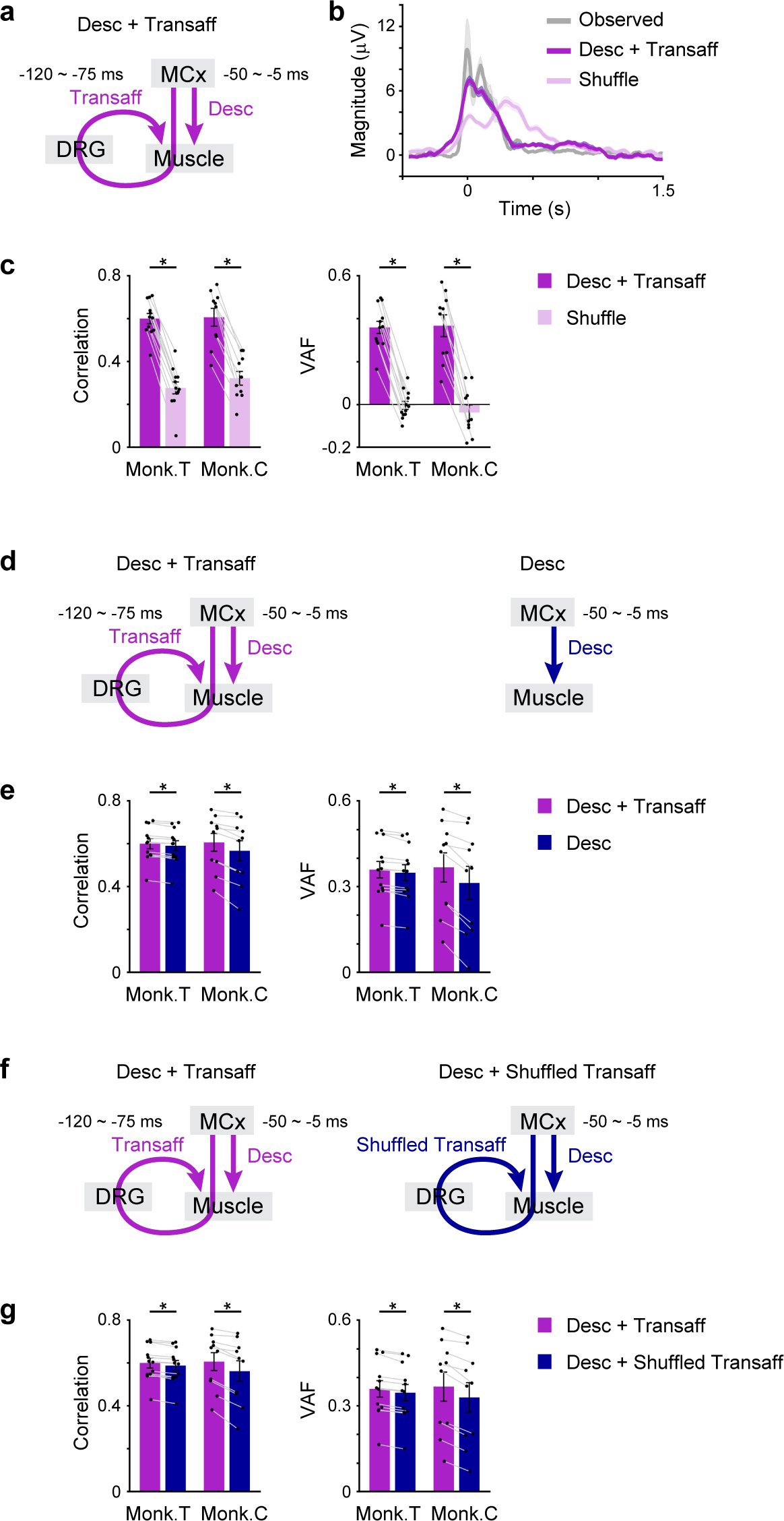
Transafferent input is important for the reconstruction of muscle activity. **a**, Model accounting for muscle activity evoked by descending and transafferent inputs. **b**, The observed muscle activity, its reconstruction using descending and transafferent inputs, and shuffled control data aligned to movement onset. The shaded areas indicate the SEM. **c**, Mean reconstruction accuracies of the models obtained from descending and transafferent inputs (purple) and from descending input only (dark blue). Correlation coefficients and VAFs between the observed and reconstructed traces (monkey T, 12 muscles; monkey C, 10 muscles). **d**, Models accounting for muscle activity evoked by descending and transafferent inputs or descending input alone. **e**, Mean reconstruction accuracies of the models obtained from descending and transafferent inputs (purple) and from descending input only (dark blue). Correlation coefficients and VAFs between the observed and reconstructed traces (monkey T, 12 muscles; monkey C, 10 muscles). **f** and **g**, Same as in **d** and **e** for models obtained from descending and transafferent inputs (purple) and from descending and shuffled transafferent inputs (dark blue). In **c**, **e**, and **g**, data are the mean ± SEM. \**P* < 0.05, paired two-tailed *t*-test.

**Extended Data Fig. 3.**
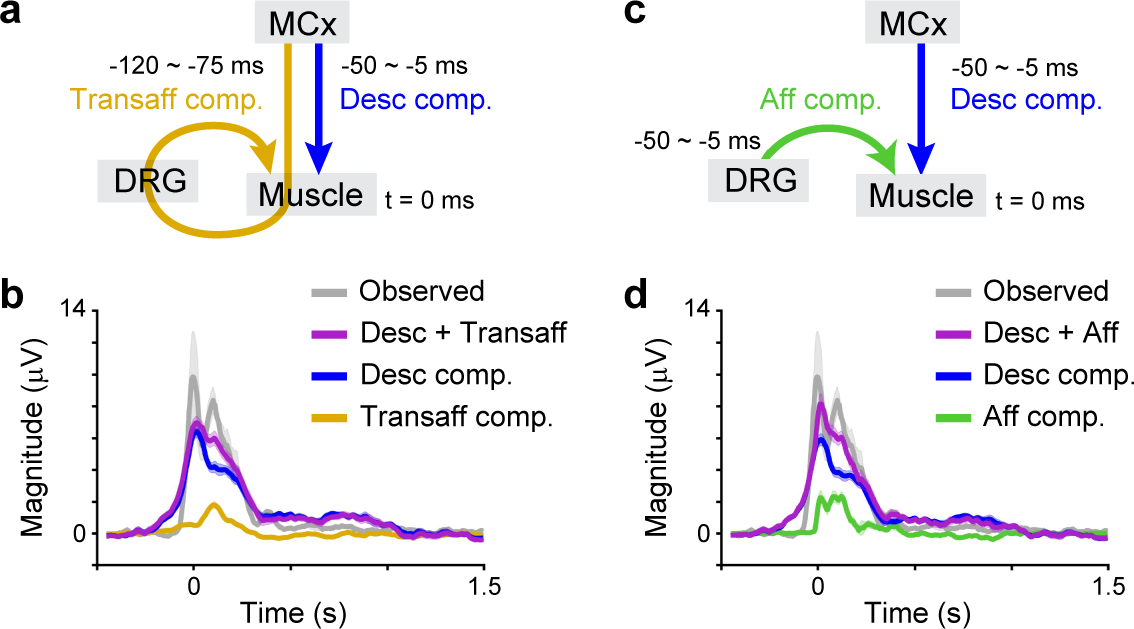
Multivariate information decomposition elucidates subcomponents. **a**, Models accounting for muscle activity evoked by descending and transafferent inputs. **b**, The observed muscle activity, its reconstruction using descending and transafferent inputs, and each subcomponent aligned to movement onset. The shaded areas indicate the SEM. **c** and **d**, Same as in **a** and **b** for descending and afferent inputs.

**Extended Data Fig. 4.**
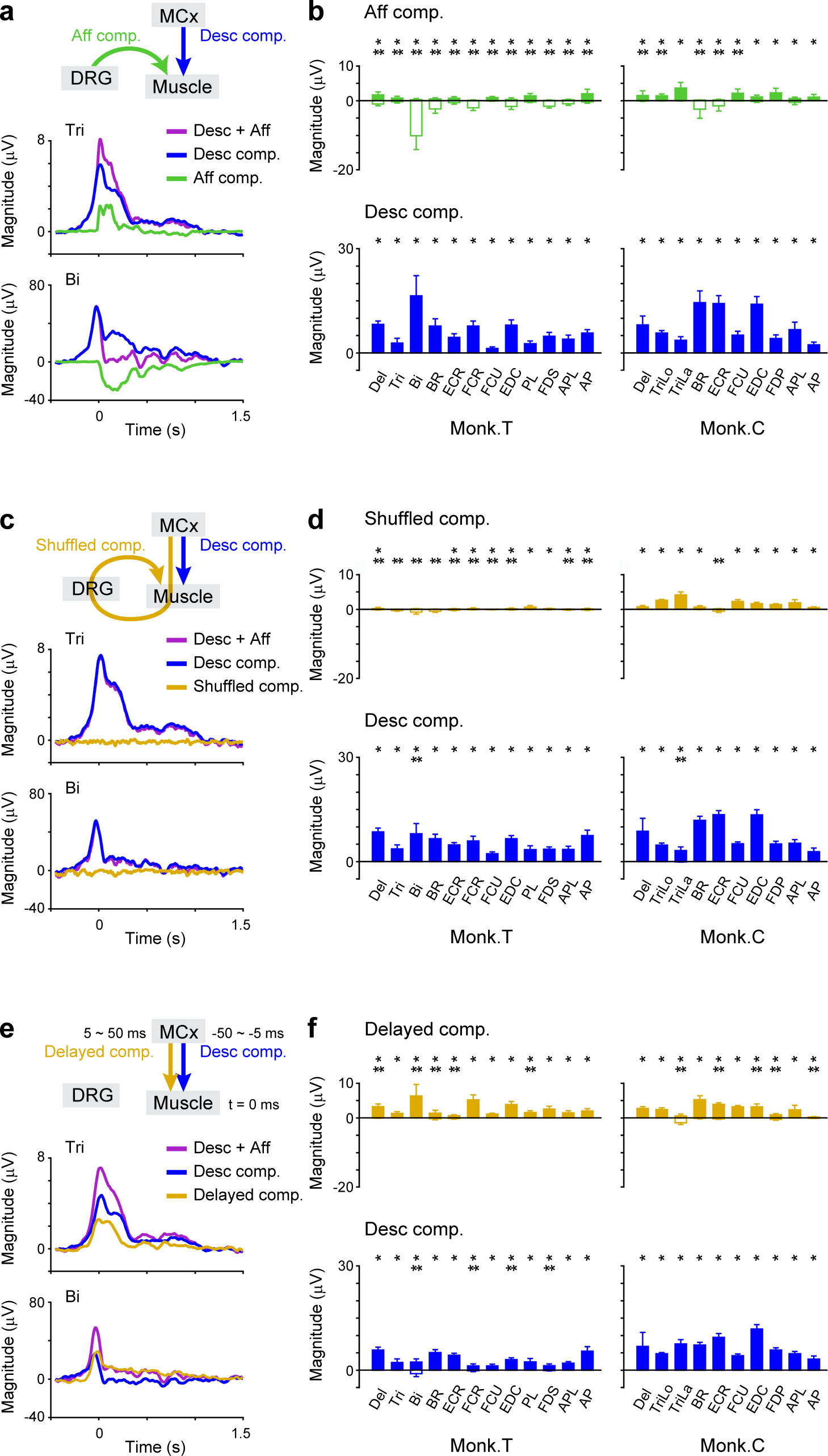
Control models fail to duplicate the corresponding component obtained by a model that accounts for descending and afferent inputs. **a**, Reconstruction of the activity of a shoulder muscle (triceps) and wrist muscle (FCR) using descending and afferent inputs, and each component in the reconstruction aligned to movement onset. **b**, Modulation of descending and afferent components for each muscle during muscle activity (monkey T, 21 sessions; monkey C, 7 sessions). **c** and **d**, Same as in **a** and **b** for descending inputs and shuffled data from transafferent inputs. **e** and **f**, Same as in **a** and **b** for descending inputs and delayed activity in the MCx. In **b**, **d**, and **f**, data are the mean ± SD. * and \*\**P* < 0.05, unpaired two-tailed *t*-test for positive and negative values.

**Extended Data Fig. 5.**
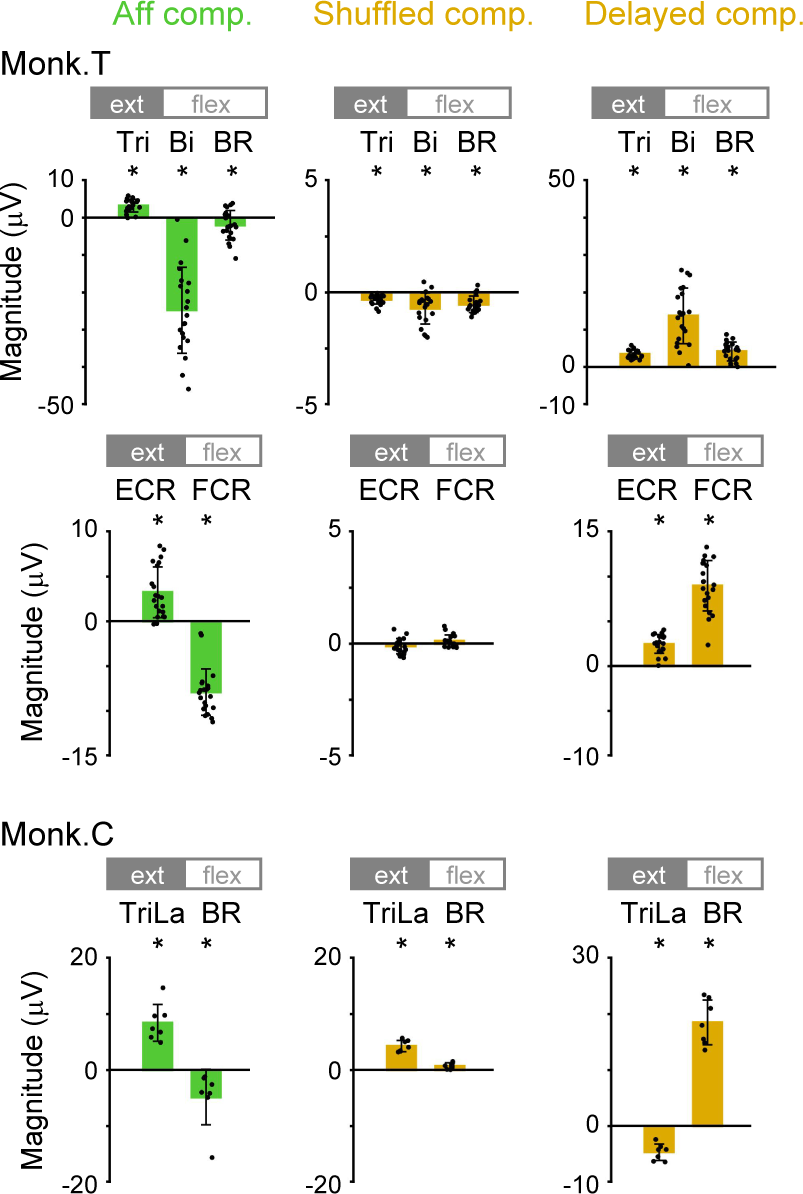
The relationship between corresponding components in control models and the initial movement is not consistent with the spinal reflex. Size of the afferent (left row), shuffled (middle row) and delayed (right row) components for antagonistic muscle pairs (ext: extensor; flex: flexor) in a period from the beginning of the reaching movement (55 to 100 ms around movement onset; shown in the green and orange areas in Fig. 4b). Asterisks indicate a significant difference from 0 (monkey T, 21 sessions; monkey C, 7 sessions). Data are the mean ± SD. \**P* < 0.05, unpaired two-tailed *t*-test.

**Extended Data Fig. 6.**
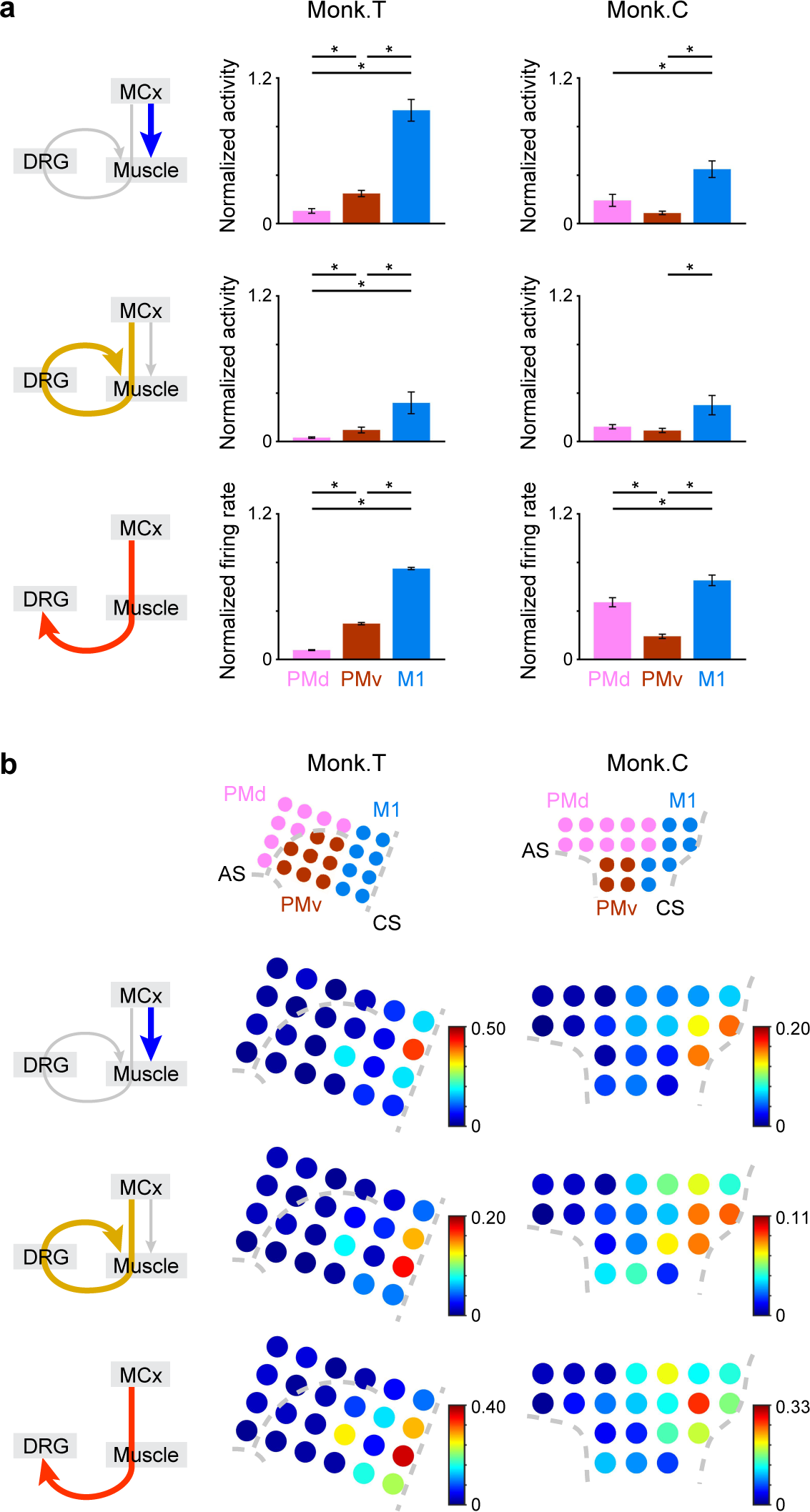
M1 is a major predictor of muscle activity by descending and transafferent pathways. **a**, Size of subcomponents calculated from the activity in each cortical area for the prediction of muscle or afferent activity (monkey T, 12 muscles and 702 afferents; monkey C, 10 muscles and 92 afferents). The thick, colored arrows in the left-hand schematics show the inputs for which the modulation is represented. **b**, Color maps in the lower row represent the size of subcomponents calculated from the activity at each electrode for the prediction of muscle or afferent activity. The position of electrodes in each motor cortical area is depicted in the upper row. The size of each subcomponent is normalized by the size of the reconstruction of muscle activity using descending and transafferent inputs or the reconstruction of afferent activity using descending input. CS, central sulcus; AS, arcuate sulcus. The thick, colored arrows in the left-hand schematics show the inputs for which the modulation is represented. In **a**, data are the mean ± SEM. *P* < 0.05, one-way repeated-measures ANOVA, \**P* < 0.05, paired two-tailed *t*-test.

**Extended Data Fig. 7.**
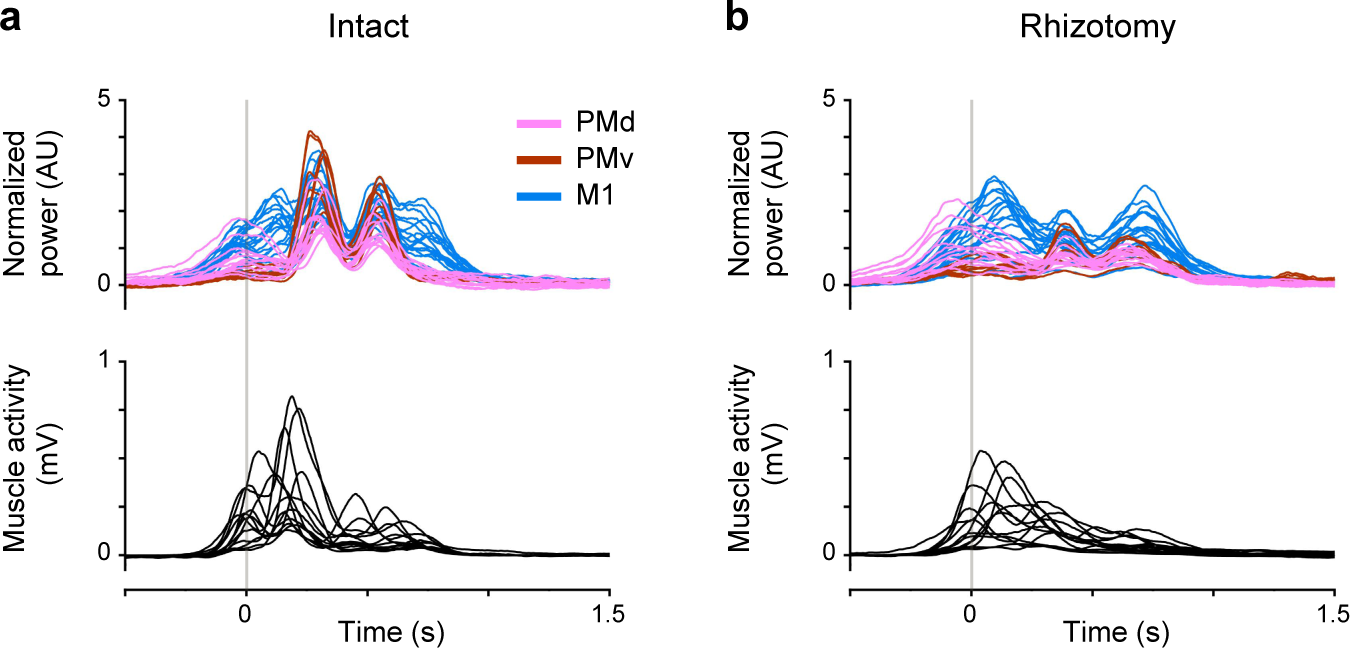
Simultaneous recording of MCx and muscle activity before and after dorsal rhizotomy. **a**, Modulation of cortical and muscle activity in monkey P aligned to movement onset before dorsal rhizotomy. *Top*: High-gamma cortical activity. *Bottom*: Forelimb muscles. A vertical line represents the time of movement onset. **b**, Same as in **a**, but after dorsal rhizotomy.

**Extended Data Fig. 8.**
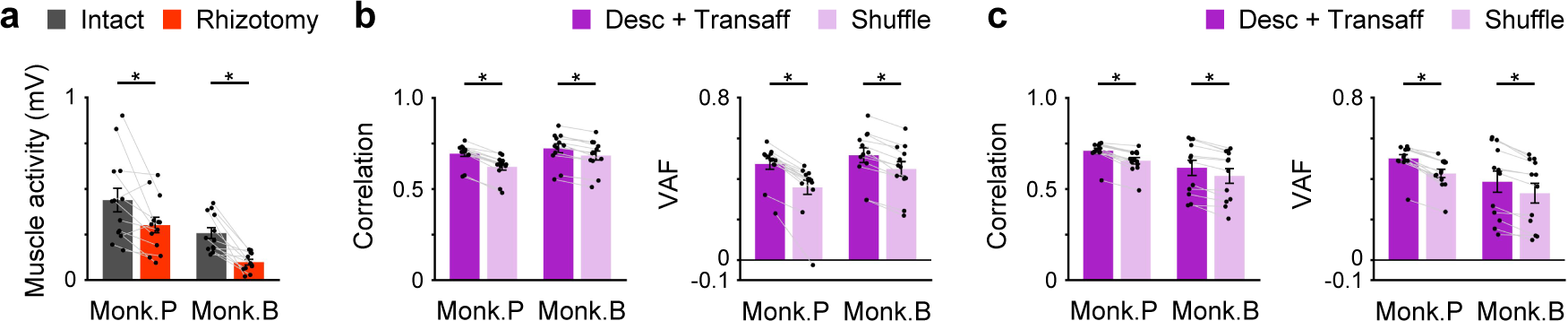
Dorsal rhizotomy decreases the transafferent effect of the MCx on muscle activity. **a**, Mean peak amplitude of muscle activity was compared before and after dorsal rhizotomy (monkey P, 13 muscles; monkey B, 12 muscles). **b**, Correlation coefficients and VAFs between the observed and reconstructed traces before dorsal rhizotomy (monkey P, 13 muscles; monkey B, 12 muscles). **c**, Same as in **b**, but after dorsal rhizotomy. In **a** to **c**, data are the mean ± SEM. \**P* < 0.05, paired two-tailed *t*-test.

**Extended Data Fig. 9.**
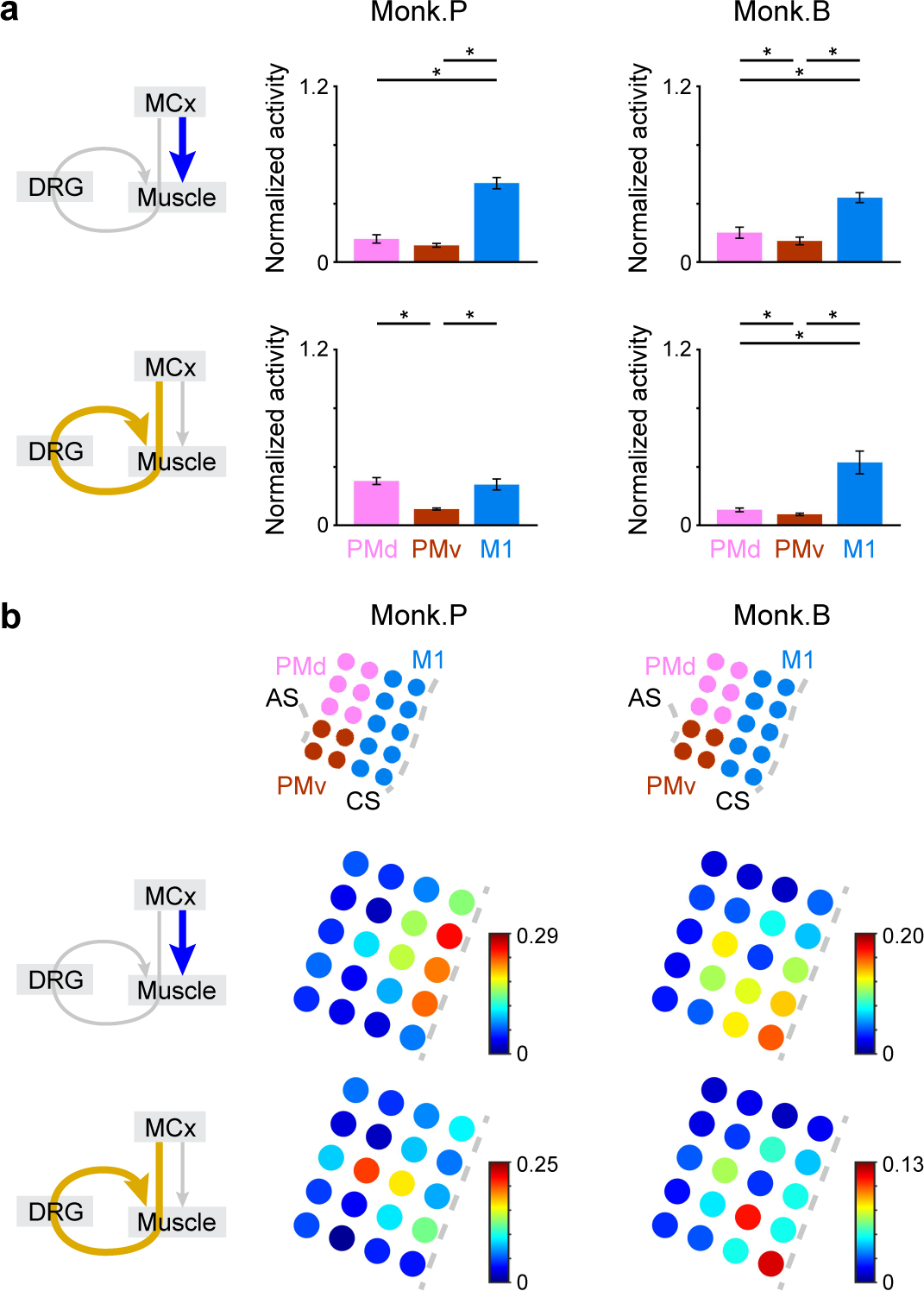
M1 is a major predictor of muscle activity by descending and transafferent pathways. **a**, Size of subcomponents calculated from the activity in each cortical area for the prediction of muscle (monkey P, 13 muscles; monkey B, 12 muscles). The thick, colored arrows in the left-hand schematics show the inputs for which the modulation is represented. **b**, Color maps in the lower row represent the size of subcomponents calculated from the activity at each electrode for the prediction of muscle. The position of electrodes in each motor cortical area is depicted in the upper row. The size of each subcomponent is normalized by the size of the reconstruction of muscle activity using descending and transafferent inputs using descending input. CS, central sulcus; AS, arcuate sulcus. The thick, colored arrows in the left-hand schematics show the inputs for which the modulation is represented. In **a**, data are the mean ± SEM. *P* < 0.05, one-way repeated-measures ANOVA, \**P* < 0.05, paired two-tailed *t*-test.

**Extended Data Fig. 10.**
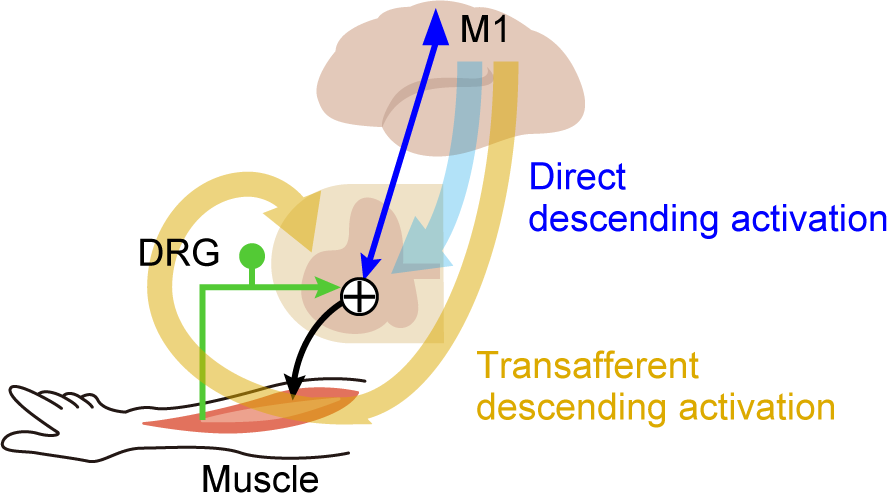
Schematic diagram illustrating parallel signal transmission from the M1 to muscles during voluntary movements. The M1 drives muscle activity via not only direct descending activation through the descending pathway but also transafferent descending activation through the transafferent pathway composed of the descending and spinal reflex pathways during voluntary forelimb movements. The M1 implements an internal model that prospectively estimates muscle activity through the spinal reflex.

## Notes

### Competing Interest Statement

The authors have declared no competing interest.

